# Red Fluorescent Carbon Nanoparticles Derived from *Spinacia oleracea L*.: A Versatile Tool for Bioimaging & Biomedical Applications

**DOI:** 10.1101/2023.05.09.540029

**Authors:** Ketki Barve, Udisha Singh, Krupa Kansara, Payal Vaswani, Pankaj Yadav, Ashutosh Kumar, Dhiraj Bhatia

## Abstract

Carbon-based fluorescent quantum dots are an emerging class of nanoparticles for targeted bioimaging and biomedical applications. We present a facile microwave-assisted approach for synthesizing carbon nanoparticles with bright red fluorescence using ethanolic extracts of *Spinacia oleracea* leaves, with a quantum yield of 94.67%. These nanoparticles, called CNPs, ranging from 15-50 nm, demonstrated fluorescence emission in the near-infrared (NIR) region between 650 and 700 nm, independent of excitation wavelength. Upon excitation at a wavelength of 410 nm, they exhibit an emission maxima peak at 672 nm. The significant uptake of CNPs in mammalian cells and zebrafish larvae highlights their potential as bioimaging agents in diverse biomedical applications *in vivo*. Further, these quantum dots enhance cellular proliferation and migration as observed by wound healing assay in mammalian cells, indicating their possible application in tissue engineering and regenerative medicine. These findings suggest that biosynthesized carbon nanoparticles possess significant potential for biomedical activities, which can serve as a robust benchmark for researchers towards promoting sustainability.

## 1. Introduction

Nanoparticles have garnered much interest in recent years with their distinctive physical and chemical characteristics. Carbon nanoparticles, in particular, have been of great interest in various fields, including targeted therapeutics, biosensing, theranostics, bioimaging, and photodynamic therapy (PDT), owing to their biocompatibility, low toxicity, and high stability. Carbon-based nanoparticles possess distinct optical properties and surface modifications due to their unique size and structure. They can be used as contrasting agents for imaging biological entities such as cells and tissues. Their small size and biocompatibility also make them suitable for targeted delivery of drugs to specific cells or tissues. The optical characteristics of CNPs are dependent on the band gap energy. Upon absorption of light at a particular wavelength corresponding to the band gap energy, the valence band electrons of these nanoparticles undergo a quick transition to the conduction band and return to the valence band by releasing energy in the form of light or photons. Compared to chemical substances (such as citric acid, glucose, urea, and thiourea) used to make CNPs, the synthesis of CNPs using natural materials has many benefits since they are less expensive, non-toxic, and more abundant.^1^ Consequently, there is a need to investigate easily accessible and cost-effective carbon sources using environmentally friendly green technologies to synthesize high-quality carbon-based nanoparticles with superior photoluminescence (PL) properties.^2,3,4^

In this regard, spinach extract comprises bio-reductants, including flavonoids, polyphenols, amino acids, and carbohydrates, that stabilize nanoparticle synthesis and mitigate agglomeration.^5^ Its application in medicine involves injecting the body to treat specific cancer cells via external magnetic field guidance.^6^ These spinach-derived nanoparticles are non-toxic and can be utilized for medicinal purposes in the pharmaceutical industry. Such a green synthesis of these nanoparticles represents an economically feasible production method.^7^ An earlier study by Xu et al.^2^ has already demonstrated the synthesis of carbon-based nanoparticles using spinach and polyethyleneimine, resulting in red fluorescence. However, those nanoparticles exhibited instability, gradually shifting from red to blue fluorescence within a week. To fabricate nanoparticles that exhibit red fluorescence over several months, we synthesized carbon nanoparticles via a simple and fast approach using the microwave. Red fluorescent nanoparticles, especially under NIR excitation (800-1200 nm), are more advantageous than blue and green fluorescent nanoparticles because they can penetrate deep tissue with minimal auto-fluorescence and a high signal-to-noise ratio in biological tissues and fluids. Also, their requirement for a low excitation energy source does not damage cells or tissues, as seen in blue fluorescence. Blue fluorescence requires high-energy excitation, which can cause cell damage and cancer. Also, autofluorescence may occur as certain plant and animal tissues exhibit fluorescence when exposed to ultraviolet (UV) radiation, leading to interference with the signal of other CNPs.^8.9^ Having said this, however, it remains difficult to synthesize red emissive CNPs with high quantum yield (QY) because the sp2 π-conjugation domains are much larger, making them more prone to deformities and vulnerable to environmental disruption.^9,10,11^

We have synthesized novel red fluorescent carbon nanoparticles by microwave synthesis route, as it combines the speed and uniform heating of the materials. The main benefit of employing microwave heating is the uniform, non-contact heat transfer to the solution, which has penetration characteristics and makes it possible for reactions to occur swiftly.^12^ The microwave method is a highly effective procedure for nanoparticle synthesis, owing to its numerous benefits, such as being non-toxic, straightforward, scalable, and cost-effective. Nevertheless, the primary drawback of this technique is the limited ability to control the size of the resulting nanoparticles precisely.^13^ Majority of synthesis methods involve temperatures of approximately 150°C or higher, primarily through hydrothermal or reflux methods. Furthermore, these methods typically require a minimum reaction time of 60 minutes or more, whereas implementing microwave-assisted approaches enables a significantly shorter reaction time of less than 30 minutes.

The carbon nanoparticles synthesized by this approach have led to a very high quantum yield of 94.67%, with high-intensity fluorescence in the near Infrared region. Further characterization revealed that they are crystalline, showing good photostability. The emission is independent of excitation wavelength, and the emission maxima are obtained at 672 nm when excited at 410 nm wavelength. To utilize their nano-size, fluorescence property, and biocompatibility, the uptake of CNPs was studied as a bioimaging agent in human cell line Retinal Pigment Epithelium cells (RPE-1 cells) and in zebrafish larvae (**Scheme 1**). This study revealed that they are effective bioimaging agents showing bright red fluorescence inside cells and larvae. As reported earlier^14^, CNPs help in the wound-closing process, so a scratch assay was performed to investigate the effect on cell proliferation. The results showed that the wound closure rate in the control group was 40%, while it was 85% in the group treated with CNPs. Increasing the concentration of CNPs also accelerated the wound-closure process. These findings indicate that the CNPs synthesized in this study have the potential as a theranostic agent and can be further modified for future biomedical applications.

**Scheme 1:**
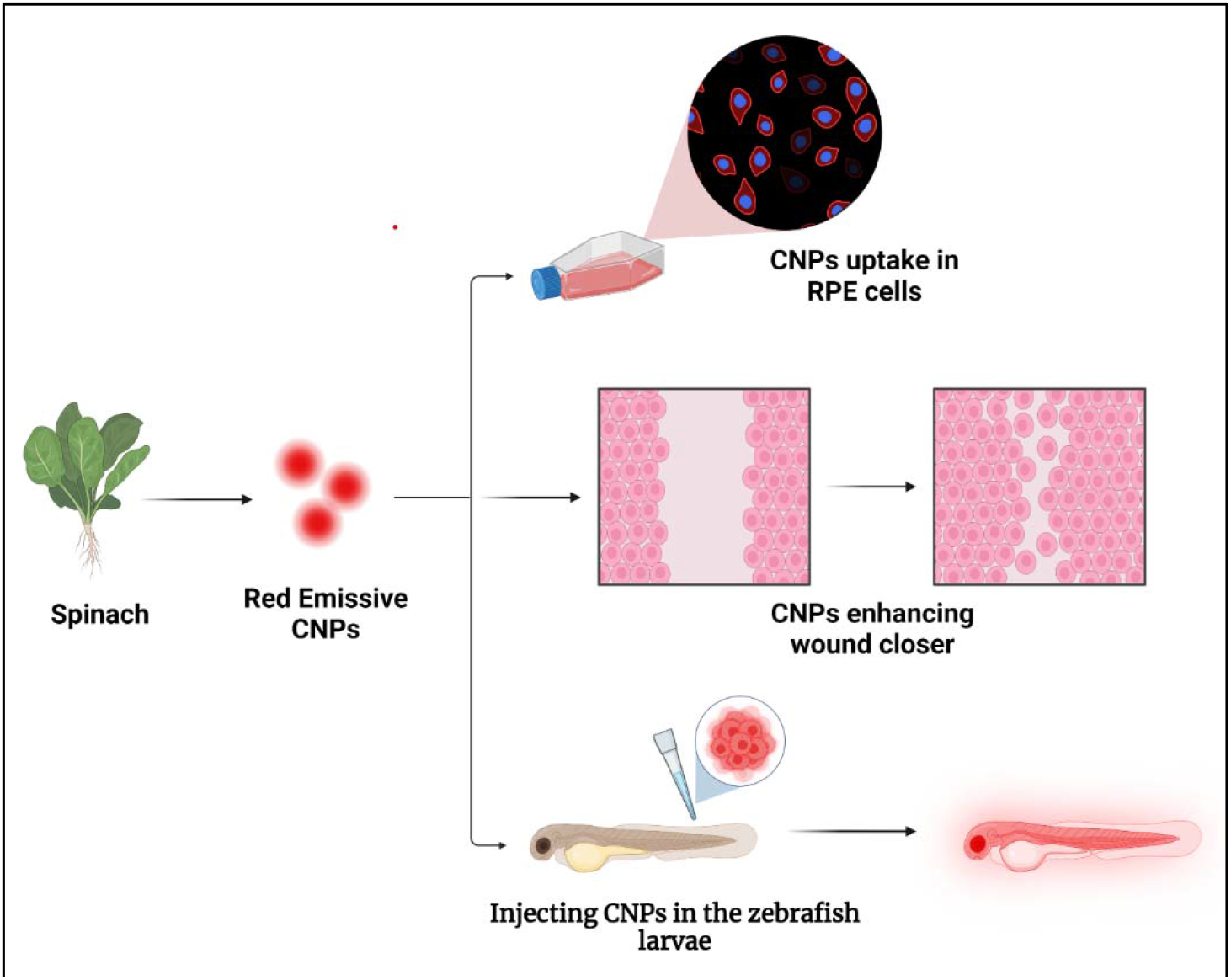
Red emissive carbon nanoparticles (CNPs) synthesized from spinach (plant) source, demonstrating potential applications in bioimaging retinal pigment epithelium cells and zebrafish larvae. In addition, they have been shown to enhance wound closure in scratch assay experiments.

## 2 Results and discussion

### 2.1 Synthesis and Characterization of CNPs

Carbon nanoparticles emitting red fluorescence were synthesized through microwave-assisted “green synthesis” approach from ethanolic extract of spinach powder. 2g of dried spinach leaf powder was mixed in 20 mL ethanol solution and kept for stirring for 4 hours on a magnetic stirrer. The resulting mixture was centrifuged at 24°C for 10 minutes at 10,000 rpm. A rota-evaporator was used to evaporate the ethanol from the supernatant. The resulting slurry was re-dispersed in 5 ml Milli-q water. The solution was microwaved until the water evaporated. After cooling, 5ml ethanol was added, properly mixed, and set aside for 30 minutes. The solution was then probe-sonicated then filtered using a 0.22 μm syringe filter. The solution was subsequently utilized to carry out further characterization and experimental research. To investigate the optical features of CNPs, we measured the absorption using UV-Vis (Ultraviolet-Visible) absorbance spectroscopy, which revealed a distinctive peak at 418 nm and 672 nm, which corresponds to the level of energy transition associated with the angstrom-scale conjugated π-structure14 as depicted in **Figure 1 (a)**. Several excitation wavelengths were used to determine the fluorescence emission spectra of red-emissive CNPs. As observed in **Figure 1(b)** the highest emission spectra (λmax) was observed at 672 nm when excited at 410 nm wavelength. To find out more about the structure and composition of CNPs, we carried out X-ray diffraction (XRD) and Fourier Transform Infrared (FT-IR) spectroscopy. X-Ray diffraction (XRD) spectra show they are crystalline. The XRD profile of CNPs indicates a single at 31.7° correspond, revealing the CNPs to be crystalline as shown in **Figure 1(c)**. Fourier transforms infrared (FTIR) spectra of CNPs carried out to detect the functional groups and chemical composition, as shown in **Figure 1(d)**. Chemical bonds between N, H, C, and O elements are usually formed on carbon-based nanoparticles. We obtained N-O stretching (1539.02 cm-1), C-H bending (880.00 cm-1), C-H stretching (2971.86, 2875.77 cm-1), and C-O stretching (1045.81, 1085.16 cm-1). We got the absorption peak at (1539.02 cm-1) that confirms the presence of Nitric oxide (N-O) stretching, (1045.81, 1085.16 cm-1) means (C-O stretch) ensures the carbonyl group is present, (880.00 cm-1) confirms that presence of C-H Bending, (2971.86, 2875.77 cm-1) affirms that the alkyne group is present on the surface of CNPs. The atomic force microscopy (AFM) image indicates a topological height of 3.14 nm, as shown in **Figure 1 (e)**, and in **Figure 1 (f)** the size of CNPs is shown approximately 50 nm. The CNPs with a production yield of 28 mg/ml and a quantum yield of 94.67% were obtained from an ethanolic extract of spinach powder (2 g).

**Figure 1.**
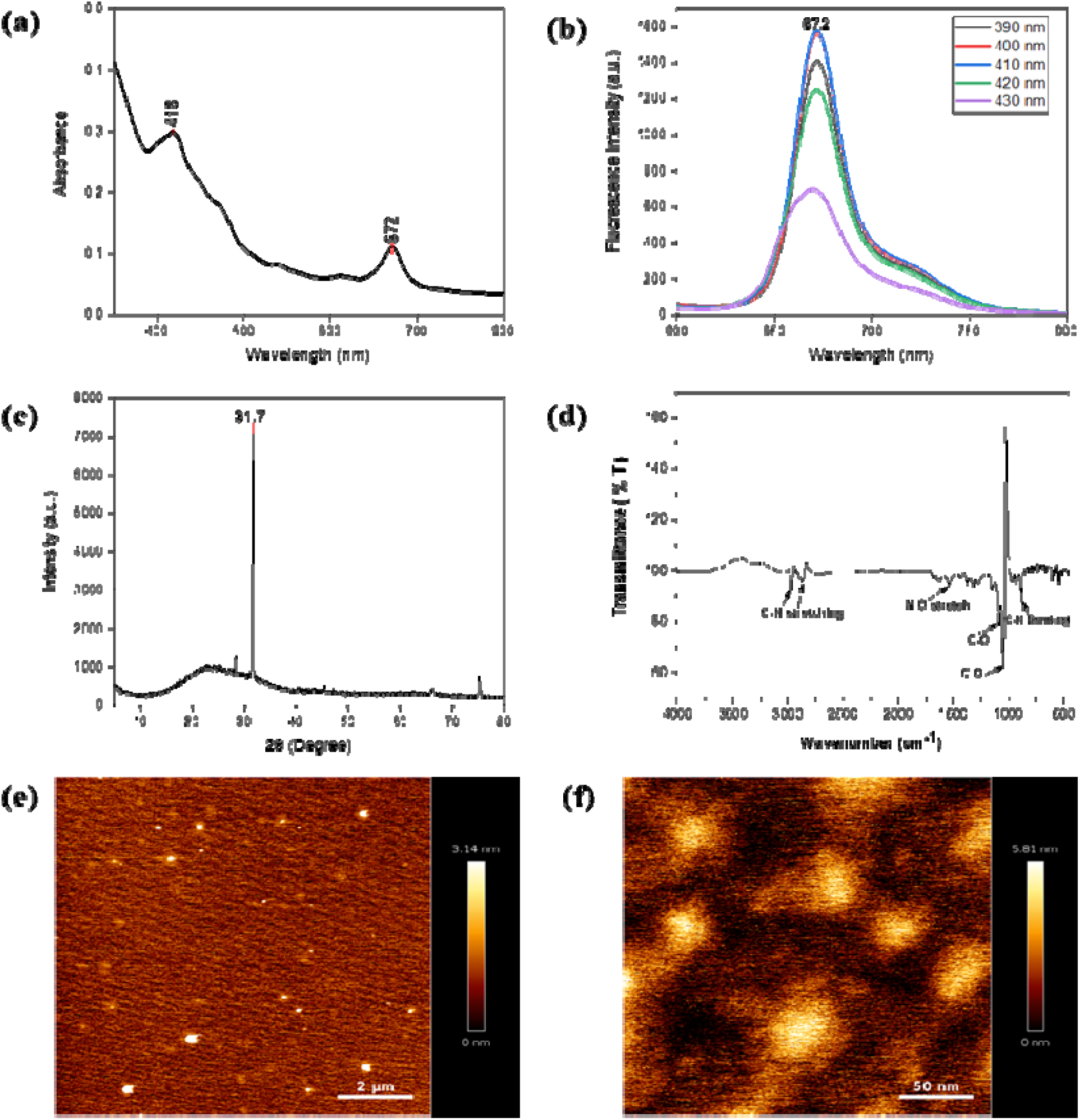
Characterization of CNPs. (a) Ultraviolet (UV)-Visible spectra of CNPs depicting two peaks at 418 nm and 672 nm related to different levels of electronic transition. (b) The excitation and emission spectra of CNPs at 410 nm excitation wavelength showed λmax at 672 nm. (c) X-Ray diffraction (XRD) analysis shows they are crystalline. The XRD profile of CNPs shows a single at 31.7° correspond, revealing the CNPs to be crystalline. (d) FTIR spectra reveal the chemical composition and functional groups of CNPs. (e) AFM of prepared CNPs using the microwave-assisted green synthesis route showed the particle size to be approximately 50 nm with a topological height of 3.14 nm. (f) Magnified image of AFM image of CNPs.

### 2.2 Optical Properties of CNPs

We evaluated the UV-VIS absorption and fluorescence emission spectra to study the optical behavior. The prepared stock solution of 28mg/ml was further diluted in ethanol for further characterization. The fluorescence intensity is 1572 a.u. at 672 nm, as depicted in **Figure 1(b)**. The emission wavelength falls between 630 nm to 750 nm range. The CNPs showed green color under daylight and bright red color under UV light. The CNPs exhibit an excitation wavelength-independent fluorescence spectrum profile. There was no apparent change in the emission profiles as the excitation wavelength increased. The QY is determined as the ratio of emitted photons to absorbed photons, which reflects the proportion of excited molecules that have returned to the ground state by releasing emission photons. The fluorescence emission experiments were carried out with Jasco spectrofluorometer model FP-8300 (Japan) in the 410 nm excitation range. To record the emission spectra of CNPs, 10 mm path-length quartz cuvettes having 10nm slit width were used. According to the excitation spectra of CNPs, rhodamine B was chosen as the reference standard for calculating the QY of CNPs. The CNPs fluorescent QY is 94.67% in ethanol by referring to standard rhodamine B. The dispersion of the synthesized CNPs is more excellent in protic solvents, such as alcohols, due to the formation of an H-bond with a nitrogen atom and the solvent molecule. As a result, it creates an inversion of the lowest-lying n-π* state to π-π*, resulting in a rise in fluorescence QY. The fluorescence quantum yield (QY) is a crucial factor in determining the fluorescent probes used. QY is influenced by various environmental parameters such as temperature, viscosity, pH, H-bonding, solvent polarity, and quenchers, as well as fluorescence lifetime. Before utilizing CNPs as a fluorescent probe, it is essential to assess all these factors meticulously. CNPs’ thermal stability was examined by exposing them to a temperature varied from 10°C to 70°C at intervals of 10°C and evaluating changes in their fluorescence spectra via a spectrophotometer. We can see from **Figure 2(a)** that the increase in temperature caused a gradual reduction in fluorescent intensity, which can be attributed to the presence of thermal energy inducing electronic vibrations that result in the release of non-radiative energy.^15^ Furthermore, time-dependent fluorescence intensity was studied at 5-minute intervals over 30 minutes and found that the intensity remained constant over time, with the peak intensity consistent with that shown in **Figure 2(b)**. Our investigation on the effects of prolonged light exposure and temperature indicates that the CNPs synthesized exhibit good photostability and thermal stability with a slight decrement in intensity as the temperature rises.

**Figure 2:**
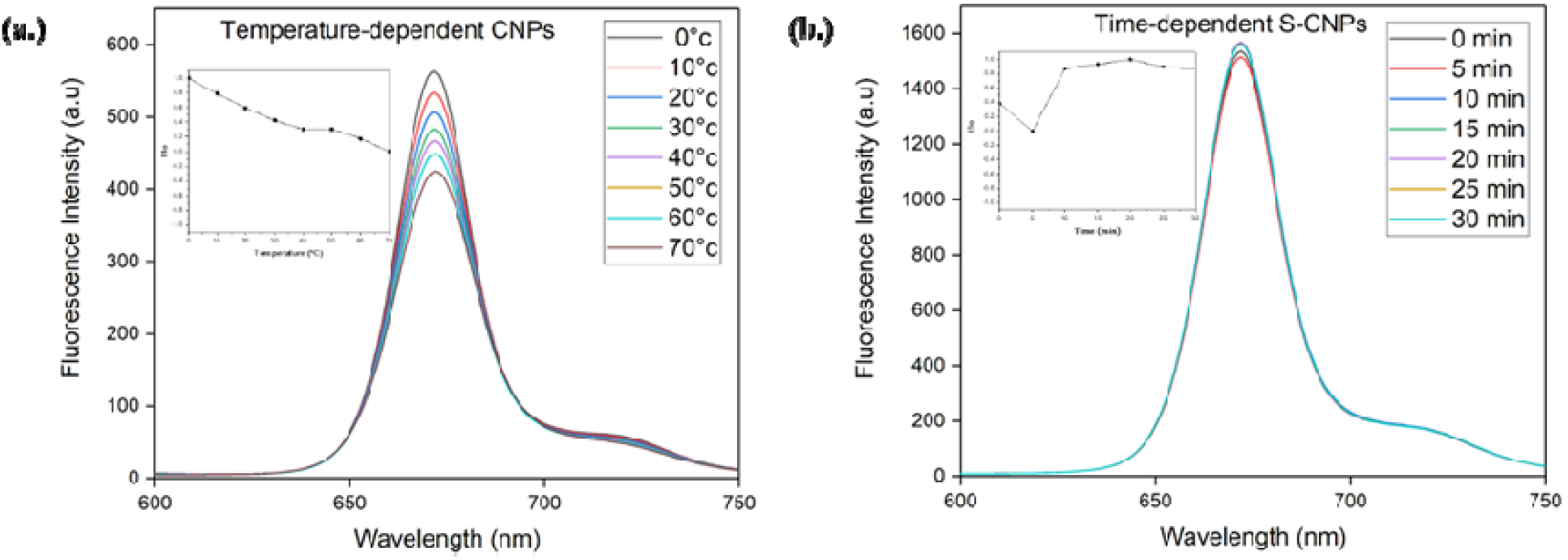
(a) At varying temperatures (0–70 °C) fluorescence spectrum of CNPs was measured with an increment of 10°C. The fluorescence intensity marginally decreases from 10°C to 70°C as the temperature rises. (b) Fluorescence spectra at different duration (0–30 min) when maintained at 27 °C. The photostability of CNP stimulated concerning time is displayed in the inset.

### 2.3 Cellular uptake studies of CNPs

Confocal microscopy analyses were carried out on retinal pigment epithelial cells cultured with CNPs at varying concentrations to investigate the potential application in bioimaging. Experiments at different CNP concentrations of 50, 100, and 200 μg/mL for 30 minutes at 37°C revealed that the CNPs were efficiently taken up by the cells and exhibited intracellular fluorescence. From inside the cells, the CNPs released a red fluorescence signal observed in the red channel and needed an excitation laser at 633 nm. As the amount of CNPs increased, the fluorescence signal was intensified, indicating a direct relationship between CNPs concentration and intracellular fluorescence intensity. Using confocal data, we analyzed and quantified the result, indicating that cellular fluorescence intensity increased with increasing CNP concentration, showing a concentration-dependent absorption and fluorescence response. Through concentration-dependent studies, it has been revealed that as CNP concentrations increase from 50 to 200 g/ml, fluorescence intensity escalates, depicted in **Figure 3**.

**Figure 3.**
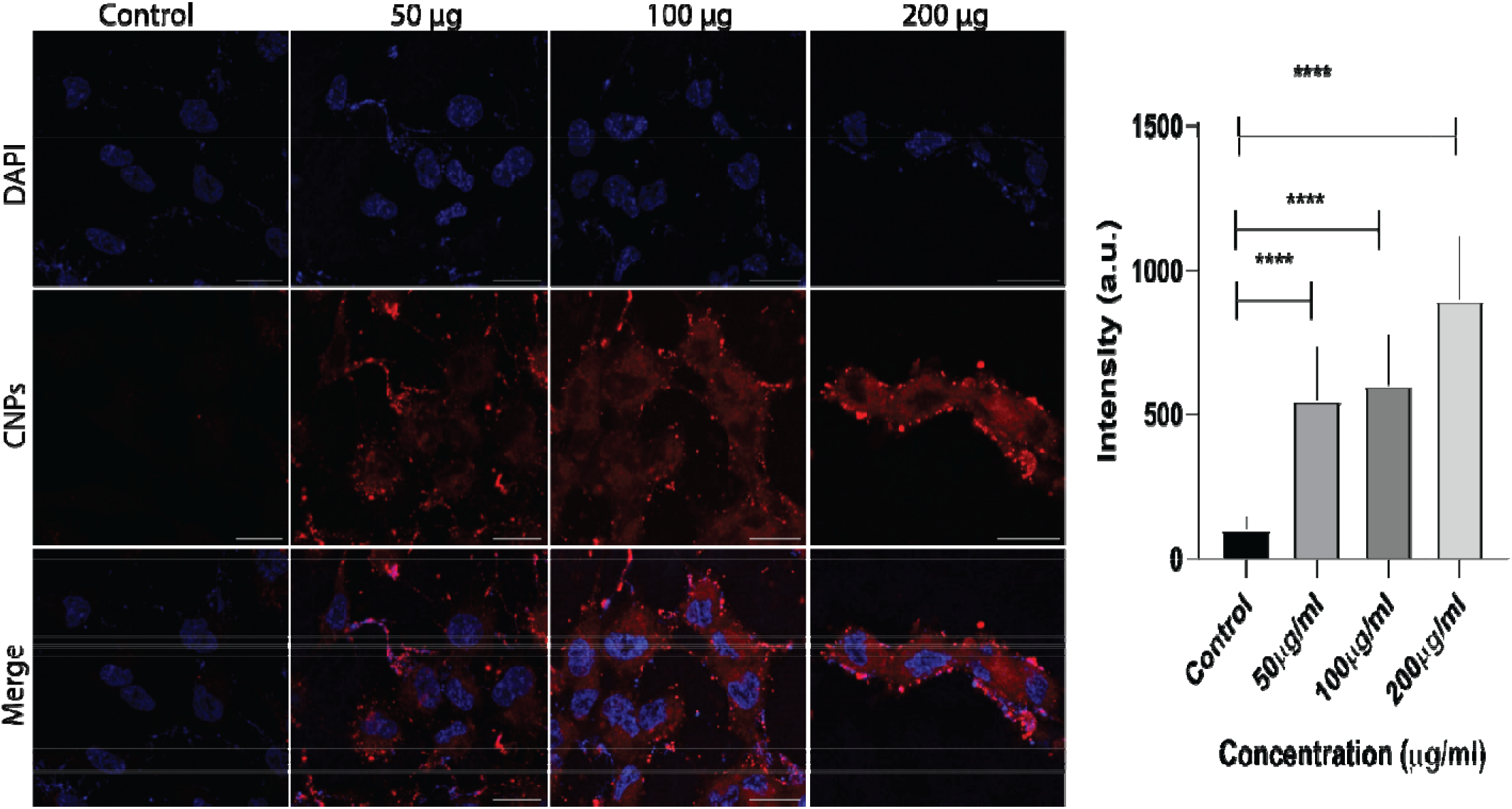
Confocal microscopy images show RPE-1 cells treated with CNPs at varying concentrations of 50, 100, and 200 μg/mL, with a corresponding control that was without CNPs, showing no fluorescence when excited at 633 nm. Studies on CNPs concentration-dependent properties were conducted, and the result showed that the fluorescence intensity rises as the concentration of CNPs increases. The confocal images scale bar is held constant at 40μm. Using Fiji ImageJ software, CNPs cellular uptake was also quantified at 50, 100, and 200 μg/mL. **** Indicates statistically significant value of p < 0.0001.

### 2.4 Scratch test to study cell proliferation

Endocytosis is the process by which different ligands and nanoparticles are taken up by cells. These ligands and nanoparticles can trigger signaling events at the cell membrane, which can result in a variety of responses, including cell migration. To assess the uptake and role of CNPs in cell migration, we carried out a scratch assay test to investigate the wound-healing abilities of CNPs. To study the effect of cell migration in wound healing, we performed the assay in RPE-1 cells. This cell line has been explored previously for analyzing migration ^5^. RPE-1 cells were cultured in six-well plates, and when the cells attained cell full confluency, a wound-mimicking scratch was created using a 200 μL tip. It was found that the CNPs didn’t hinder cell migration. To evaluate cell migration, the experiment comprised treating cells with CNPs at various concentrations and taking images at different time intervals. The findings revealed that there was an increase in cell migration and a faster wound healing response as the concentration of CNPs was increased from 50 μg/ml to 250 μg/ml, as shown in **Figure 4**. In control, where no CNPs are added, the total wound healing is around 40%, whereas, in the presence of different CNPs, the wound closure process escalates with the increase in the concentration of CNPs with almost 85% of wound healing at 250 μg/ml of concentration of CNPs.

**Figure 4.**
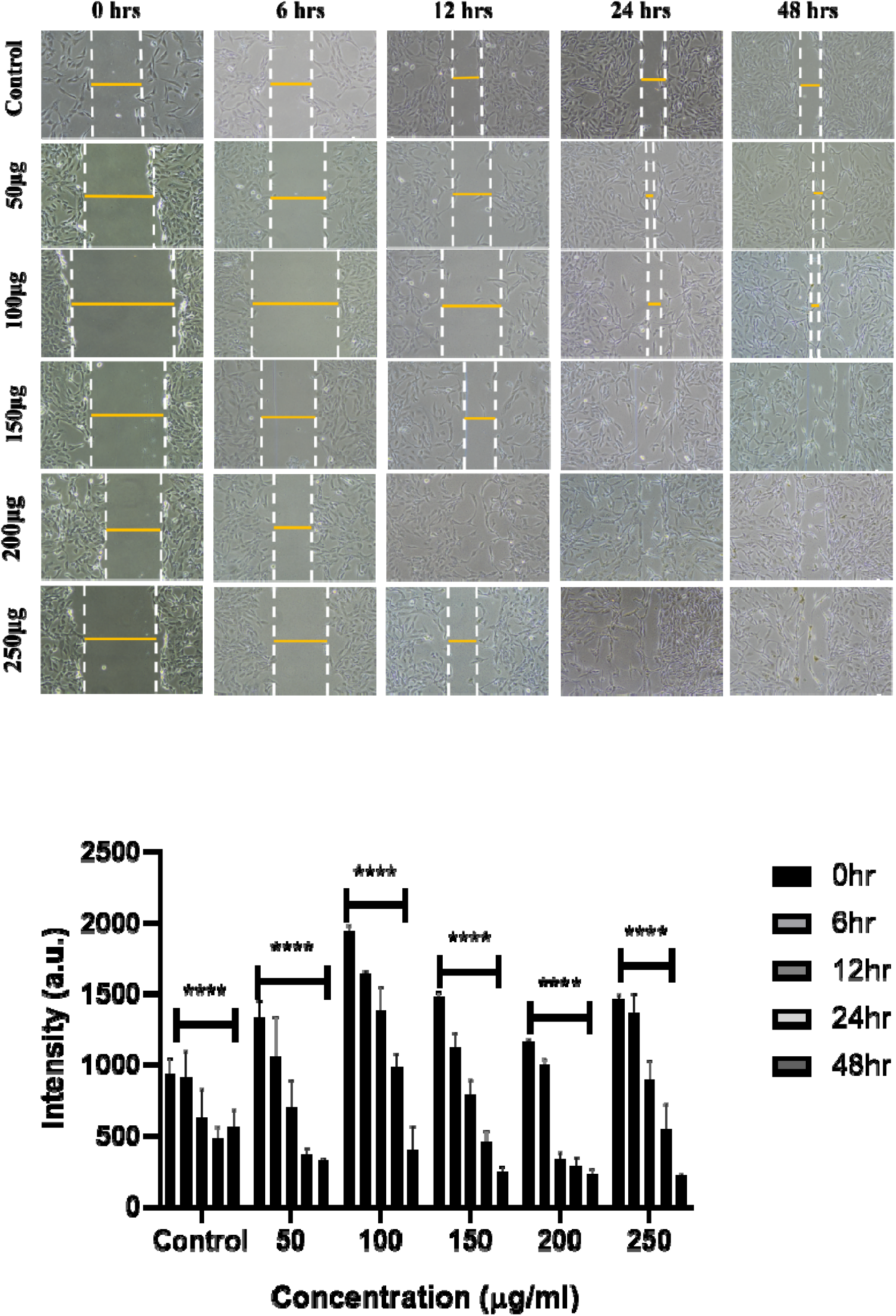
(a) The wound-healing process of RPE-1 cells was imaged at different time points using a Nikon camera. The images were captured for control and cells treated with CNPs at concentrations of 50, 100, 150, 200, and 250 μg/mL. (b) Graph depicting that the wound healing process was accelerated with increasing concentrations of CNPs compared to the control group (without CNPs). Fiji ImageJ software is being used for quantification of the wound healing process in the presence of CNPs at 50, 100, 150, 200, and 250 μg/mL concentration. **** Indicates statistically significant value of p < 0.0001, indicates statistically significant value of p = 0.002.

### 2.5. CNPs uptake in zebrafish model system

To explore the *in vivo* imaging potential of developed CNPs, we studied the uptake in the animal model, where zebrafish larvae at 72 hours post-fertilization (hpf) were administered at varying concentrations of 200 μg/ml, 300 μg/ml and 400 μg/ml of CNPs for 4 hours. The finding indicates there was an increase in the uptake of CNPs as concentration increased, which can be attributed to the improved bioavailability of these nanoformulations to the larvae. A statistically significant rise in the fluorescence intensity of the CNPs was seen at four hours post-treatment compared to the control. Furthermore, the CNPs were found to be uniformly distributed throughout the zebrafish, including the yolk sac, head, and tail regions, indicating their efficient uptake, as we can see in **Figure 5**. These findings suggest that the CNPs possess bioaccumulation properties and validate their potential for bioimaging.

**Figure 5.**
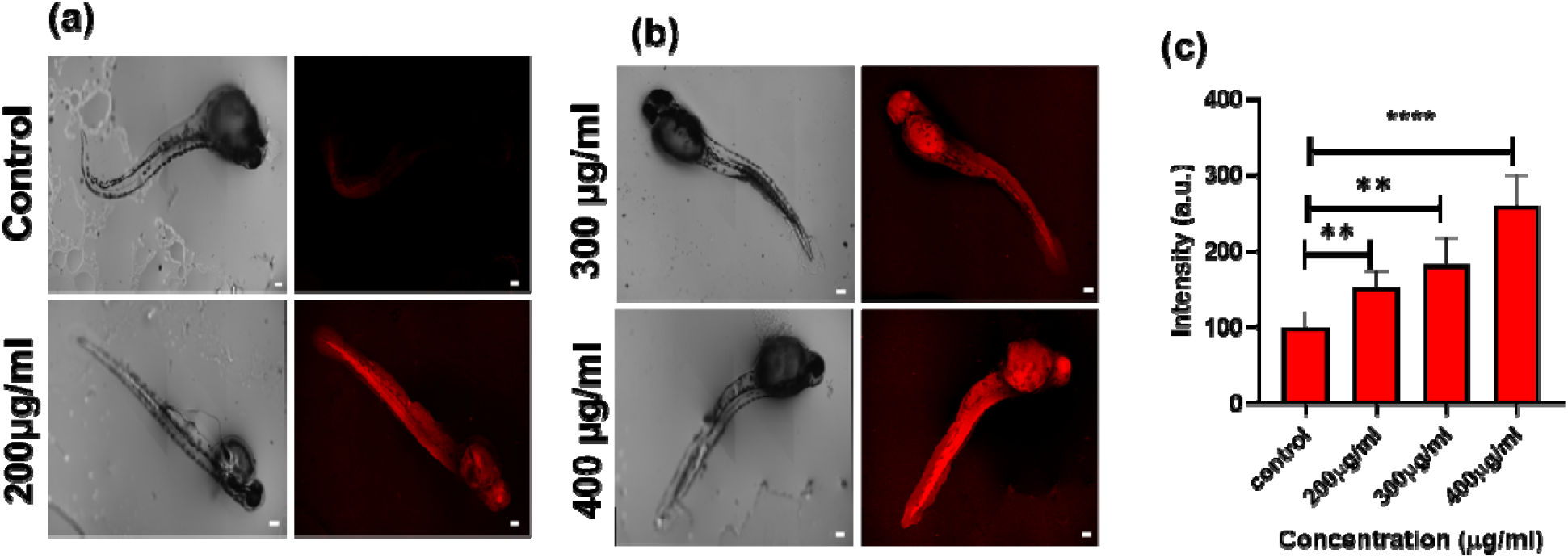
Internalization studies of CNPs in zebrafish larvae. Different concentrations (200 μg/ml, 300 μg/ml, and 400 μg/ml) of CNPs were uptaken by zebrafish larvae as shown in images (a) and (b). The zebrafish larvae were treated for 4 h with each concentration. A quantification study of the internalization of CNPs is shown in Graph (c). The scale bar is set at 100 μM. ^**^ Denotes the statistically significant p-value (p < 0.01). ^****^ denotes the significant statistical value p-value (p < 0.0001). 06 larvae were analyzed per condition.

## 3. Conclusions and discussions

Current study presents a facile and environmentally friendly method for synthesizing carbon nanoparticles using Spinacia oleracea leaf extracts. The resulting red-emissive carbon-based nanoparticles exhibit bright red luminescence in the near-infrared (NIR) region, with an emission maximum of 672 nm and a high quantum yield of 94.67%. The synthesized CNPs have demonstrated remarkable properties, had nano-size range and exhibited excellent photostability over long-term storage. The studies on CNPs also reveal their significant uptake in both RPE1 cells and zebrafish larvae, highlighting their potential as a targeted bioimaging agent in diverse biomedical applications. Scratch assay analysis demonstrates that higher concentrations of CNPs (ranging from 50μL to 250μL) led to increased cell migration and faster wound-closure response within 48 hours. Thus, it can be concluded that CNPs promoted cell migration more effectively than the control group, indicating that they are biocompatible and can accelerate wound healing, along with potentially being used as a bioimaging agent for various biological applications, including tissue engineering and regenerative medicine. Future studies will focus on maximizing their uptake efficiency and investigating their compatibility with diverse *in vivo* biological systems. Additionally, generating nanoparticles for theranostic applications combining diagnostic and therapeutic roles, could be a fruitful avenue for research. For example, conjugating drugs with nanoparticles could enable targeted drug delivery and monitoring of treatment efficacy in real time. Thus, there is a potential future for biosynthesized carbon nanoparticles for biomedical applications, which can serve as a benchmark for researchers to promote sustainability in the field of nanotechnology.

## 4. Materials & methods

### 4.1 Materials

Fresh spinach leaves were obtained from a vegetable vendor located at IIT Gandhinagar, Gujarat, India. The experiment utilized deionized water obtained from Merck Millipore, absolute ethanol (>99.9%) was supplied by Changshu Hongsheng Fine Chemicals Co., Ltd., rhodamine B was purchased from HiMedia, and syringe filters procured from Merck. Dulbecco’s modified Eagle’s medium (DMEM), penicillin-streptomycin, trypsin-EDTA (0.25%), and fetal bovine serum (FBS) were obtained from Gibco and were of excellent scientific grade, requiring no additional sterilization or treatment.

### 4.2 Synthesis of Fluorescent CNPs

The microwave approach was used to synthesize red fluorescent-emitting carbon nanoparticles. In a clean, dirt-free glass vial, 2g spinach powder was mixed in 20 mL ethanol solution and kept for 4 hours for stirring on a magnetic stirrer. The resulting mixture was centrifuged at 24°C for 10 minutes at 10,000 rpm. A rota-evaporator was used to evaporate the ethanol from the supernatant. The resulting slurry was re-dispersed in 5 ml Milli-q water. The solution was microwaved until the water evaporated. After cooling, 5ml ethanol was added, properly mixed, and set aside for 30 minutes. The answer was then probe-sonicated then filtered using a 0.22 m syringe filter. The synthesized nanoparticles dissolved in ethanol were used for further experiments.

### 4.3 Analytical methods used to Study the Characterization of CNPs

Two-dimensional and three-dimensional images of CNPs were obtained using an atomic force microscope (Bruker Multimode 8).To analyze the crystalline properties of the CNPs, A Bruker-D8 DISCOVER X-ray spectrometer, and Cu-Kα radiation were used to perform X-ray diffraction experiments. The diffraction scans were taken at a range of 5° to 80° with a scanning rate of 0.2°/min. Using a Jasco FP-8300 spectrofluorometer (Japan), the CNPs were excited in the 400–500 nm region. Rhodamine B was taken as the standard reference for calculating relative QY, and it was discovered that the CNPs had a quantum yield of 94.67% in ethanol. Using a Spectro Cord-210 Plus Analytic Jena UV-Vis spectrophotometer made in Germany, the CNP’s UV-VIS absorbance spectra were measured. FTIR of CNPs was measured in ATR (attenuated total reflectance) mode using a Spectrum 2 FTIR spectrometer from PerkinElmer. The scanning range for the measurements was 1000 cm^-1^ to 4000 cm^-1^.

### 4.4 Spectroscopic studies

The absorbance and emission spectra were used to measure using Spectroscopic-grade solvents. The emission spectra analysis was measured at 10 nm following the excitation wavelength for the standard emission spectra recording. The relative quantum yield of CNPs was calculated using Rhodamine B as a reference.

The following equation is used to calculate quantum yield:

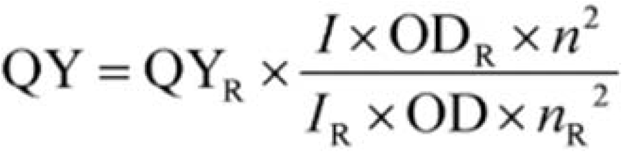

Where Quantum yield is denoted by QY, Integrated fluorescence intensity is shown as I, OD stands for optical density, n is the refractive index, and R indicates a reference.

### 4.5 Cell Culture

Professor Ludger Johannes graciously provided retinal pigment epithelial cells, RPE1 from Institut Curie in Paris, France. This cell line was cultured in complete DMEM media, and 1X PBS with a pH of 7.4 was used in all experiments.

#### 4.5.1 Confocal Microscopy Studies

RPE1 cells were seeded at a density of approximately 40,000 cells per well onto coverslips in 24-well cell culture plates containing DMEM cell culture media. The cells were then incubated for 24 hours at 37°C with 5% CO_2_. Once full confluency of RPE1 cells had reached, they were treated with PBS and then subjected to three different doses of CNPs (50, 100, and 200 μg/mL) for 30 minutes. The cells were subsequently fixed with 4% PFA around 20 minutes after being rinsed twice with PBS. After fixing, the cells were washed thrice with PBS to remove any remaining particles and PFA. DAPI stain was used to stain the fixed cells, and a mounting medium (mowiol) was used to mount them on glass slides. An oil immersion objective 63X Leica TCS SP5 confocal microscope was used to examine the cellular internalization of CNPs. A 405 nm laser was used as the excitation source for DAPI, whereas a 633 nm laser was employed for the excitation of nanoparticles. For DAPI and carbon nanoparticles, the emission bandwidths were tuned accordingly at 410–450 nm and 644–800 nm.

#### 4.5.2 Zebrafish Husbandry and Maintenance

Zebrafish of Assam wild type were brought from local vendors. They were retained in under-regulated laboratory settings at Ahmedabad University by the procedure described by Kansara et al., 2019.^16^ In artificially created freshwater, male and female fishes were kept in 20 L tanks with the quality of the water maintained by ZFIN standards. Using a multi-parameter device PCD 650 Model, Eutech, from India, water parameters such as TDS (220–320 μg/ml), pH (6.8–7.4), dissolved oxygen (>6 μg/ml), conductivity (250–350 μg/ml), and salinity (210– 310 μg/ml) were frequently examined. Zebrafish embryos were acquired and cultured for the experiment in E3 medium, composed of deionized water containing 0.33 mmol/L CaCl2, 0.17 mmol/L KCl, 0.33 mmol/L MgSO4 and 5 mmol/L NaCl, and adjusted to pH 7.2 before autoclaving the E3 media. The embryos were maintained at 28°C during the experiment. The pH of the E3 media was adjusted to 7.2, and it was autoclaved and stored at room temperature. The incubation was carried out at 28°C.

##### 4.5.2.1 In vivo Studies in Zebrafish Larvae

The uptake analyses were conducted utilizing zebrafish larvae at 72 hours post-fertilization (hpf) by the recommendations given by the Organization for Economic Cooperation and Development (OECD). Live larvae were added to six-well plates containing 15 larvae per well after dead larvae were removed. The bioimaging potential of CNPs was evaluated in larvae by administering varying concentrations (200, 300, and 400 μg/mL) of the nanoparticles in each batch. One set of control wells was kept, where larvae were not exposed to CNPs. The other set comprises varying concentrations of CNPs. After the CNPs treatment, excess CNPs were removed by washing the larvae twice with E3 media. The larvae were then placed on a glass slide via mounting Mowiol solution after being fixed using 4% PFA solution for approximately two to three minutes. The slides were then analyzed for confocal imaging after being air-dried.

## Acknowledgments

We sincerely thank all the members of the DB group for critically reading the manuscript and for their valuable feedback. KB thanks D. Y. Patil Vidhyapeeth, Pune US thanks IITGN-MHRD, GoI for Ph.D. KK thanks SERB, GoI, for the National Postdoctoral Fellowship. DB thanks SERB, GoI for Ramanujan Fellowship, IITGN for the startup grant, and DBT-EMR, Gujcost-DST & GSBTM for research grants. Imaging facilities of CIF at IIT Gandhinagar are acknowledged.

